# JM-20 administration to animals with lesion of the nigrostriatal dopamine pathway induced by 6-hydroxydopamine, partially reverses motor damage and oxidative stress

**DOI:** 10.1101/2024.07.30.605870

**Authors:** Luis Arturo Fonseca-Fonseca, Laura Reina Taño Portuondo, Jeney Ramírez-Sánchez, Nancy Pavón Fuentes, Abel Mondelo Rodríguez, Víctor Diogenes Amaral da Silva, Silvia Lima Costa, Yanier Núñez-Figueredo

**Author notes:** **Corresponding author.** Centro de Investigación y Desarrollo de Medicamentos, Ave 26, No. 1605 Boyeros y Puentes Grandes, CP 10600, La Habana, Cuba., *e-mail address* (Yanier Núñez-Figueredo).

## Abstract

Previous studies have shown that JM-20, a new chemical hybrid molecule, protects against rotenone and 6-hydroxydopamine neurotoxicity. Also, we demonstrated that JM-20 blocks the formation of toxic alpha-synuclein aggregated species and aminochrome cytotoxicity. The present study sought to determine the neuroprotective property of JM-20 in animals with a partial lesion of the nigrostriatal dopamine pathway induced by 6-hydroxydopamine (6-OHDA). For *in vivo* studies, adult male Wistar rats were lesioned in the right *substantia nigra pars compacta* (SNpc) with 6-OHDA administration. Fifteen days after surgery, the animals asymmetry levels were assessed. Those with asymmetry values higher than 50% were divided into two groups: animals that did not receive any treatment and those that were administered with JM-20 (40 mg/kg, intragastric via gavage) for 27 days. Every seven days, the asymmetry values of the animals were analyzed until day 42 after the surgery. At the end of the experiment, the animals were euthanized and the SNpc and striatum were taken out for the analysis of oxidative stress. Our results reveal a behavioral function progressively recovered in the JM-20-treated animals, diminishing the percentage of motor asymmetry. Also, it improves some oxidative stress markers in the SNpc and the striatum of these animals. Our study provides the preclinical evidence to support the long-term neuroprotective potential of JM-20 in 6-OHDA hemiparkinson rat model, pointing out to its possible use as a disease-modifying agent in PD.

## 1. Introduction

Our previous works suggested that JM-20 is an adequate compound for neuroprotection in experimental models related to different neurodegenerative disorder^1^. Different *in vitro* and *in vivo* preclinical models were performed ^2-4^ and multitarget mechanisms were demonstrated, including: (I) antioxidant and protective activities in brain-isolated mitochondria; (II) modulation of the glutamatergic system with no amnesic or addictive effects; (III) protective action against a non-physiological increase in intracellular calcium concentrations and (IV) anti-inflammatory and anti-apoptotic effects ^2-9^. In addition to the neuroprotective properties described above, its protective potential has been demonstrated in models of dementia/cognitive deficit ^10,11^ and traumatic brain injury ^12^. All these common mechanisms, which are seen in most common neurodegenerative diseases, led us to assume that it could be effective in the treatment of PD. We previously demonstrated that JM-20 bears neuroprotective capacity against rotenone and 6-hydroxydopamine-induced (6-OHDA) impairment in both *in vitro* and *in vivo* models ^13,14^. Moreover, it inhibits the formation of alpha-synuclein-aggregated species ^15^. Nevertheless, the neurorestorative potential of JM-20 is not yet investigated in any neurodegenerative preclinical models. Therefore, this study sought to assess the neurorestorative property of JM-20 in *in vivo* 6-OHDA models of PD.

## 2. Methods

### 2.1. Animals details

For this work, male Wistar rats (230-250 g) were acquired from the Center for the Production of Laboratory Animals (Havana, Cuba), housed in the animal care facility for seven days before the beginning of the *in vivo* experiments. In a temperature-controlled environment (22 ± 2ºC) with a 12-hour light/dark cycle and access to a standard diet of commercial food, as well as water *ad libitum*.

The animal care, housing and the application of experimental protocols were in agreement with international guidelines (National Institutes of Health guide for the care and use of Laboratory animals (NIH Publications No. 80-23, revised 1978), the directive 2010/63/EU of the European Parliament and the Council of 22 September 2010), in addition to institutional ones, conducted according to approved protocols (Animal Care Committee from CIDEM; code PENF 04/21). All efforts were suitably made in order to minimize the number of animals used in the experiments and their suffering. For 6-OHDA injection, the animals were placed in a stereotaxic surgical frame for rodent surgery (Stoelting Instruments, Italy). Both the surgical procedure and the preparation of the 6-OHDA were conducted as we previously described ^14^.

### 2.2. Evaluation of the JM-20 effect, experimental groups

For the evaluation of the effect of JM-20 on the 6-OHDA-induced damage model, experimental subjects were randomly assigned to three experimental groups: (I) 6-OHDA vehicle-injected animals; (II) 6-OHDA lesioned animals and treated with carboxymethylcellulose (CMC) and (III) 6-OHDA lesioned animals and post-treated with JM-20 (40 mg/kg). Vehicle (0.05 % CMC solution) or JM-20 were administered orally (intragastric (i.g.), with gavage). Immediately before use, JM-20 was prepared and suspended in a 0.05 % CMC solution and administered i.g. at 40 mg/kg dose. Administrations (CMC or JM-20) began 15 days after surgery.

### 2.3. Behavioral studies in the cylinder test

The study of motor dysfunction associated with neurotoxic injury was conducted using the cylinder test, as we had previously described ^14^: performed one day before the damage and every seven days for 42 days. On day 14 after surgery, all animals that presented a percentage of asymmetry ≥ 50% in the cylinder test (inclusion/exclusion criteria) were randomized to be included in the group with JM-20 (40 mg/kg) or without treatment (CMC). In order to determine the neurorestorative property of JM-20 in the 6-OHDA model, we decided to induce the damage. After two weeks (14 days after surgery), we used a cylinder test to determine those animals that presented a percentage of asymmetry ≥ 50%, which correlates to a degeneration of dopaminergic neurons of more than 60% ^16^. We did so in order to state whether JM-20 was able to induce any behavioral changes in animals that presented *per se* considerable dopaminergic neurodegeneration ^17-19^. One day after, the administration of JM-20 or CMC to the experimental subjects started.

### 2.4. Study of oxidative stress markers

After conducting the behavioral tests, 42 days following the 6-OHDA injection, we proceeded to the extraction of the *substantia nigra pars compacta* (SNpc) and the striatum (ST). Redox status (lipid peroxidation malondialdehyde (MDA)) and total thiol content (T-SH) groups were evaluated, only in the right hemisphere, according to the protocol described by Wong-Guerra and colleagues ^20^.

### 2.5. Statistical Analysis

GraphPad Prism 8.0.2 (263) software (GraphPad Software Inc., USA) was used to analyzed significant differences between experimental groups. Data normality was determined using Shapiro–Wilk normality test. Data were expressed as mean ± standard error of the mean (SEM) and were analyzed by one-way analysis of variance (ANOVA) or two-way ANOVA (factors: treatment × time as repeated measures) followed by the Tukey’s *post hoc* test, as indicated in the figure caption. Differences were considered statistically significant at p < 0.05.

The complete data set obtained from this study is publicly available at https://zenodo.org/records/13131465.

## 3. Results

One day before the surgery, the animals presented normal values of asymmetry, with a slight predominance in using the right paw. Fourteen days after surgery, one-way repeated ANOVA showed significant effects (F (2, 39) = 24.73 24.74, p < 0.0001) between the groups (**Fig. 1A**). One-way ANOVA and Tukey’s post hoc test analysis confirmed that were difference between the asymmetry values obtained from the animals of the 6-OHDA group concerning the pre-surgery evaluation, day -1 (p<0.0001) and with respect to vehicle group on day 14 (p<0.0001) (Fig. 1A). 6-OHDA lesioned rats, presenting a high percentage of asymmetry as compared to the control group (Vehicle) (**Fig. 1A**).

**Figure 1.**
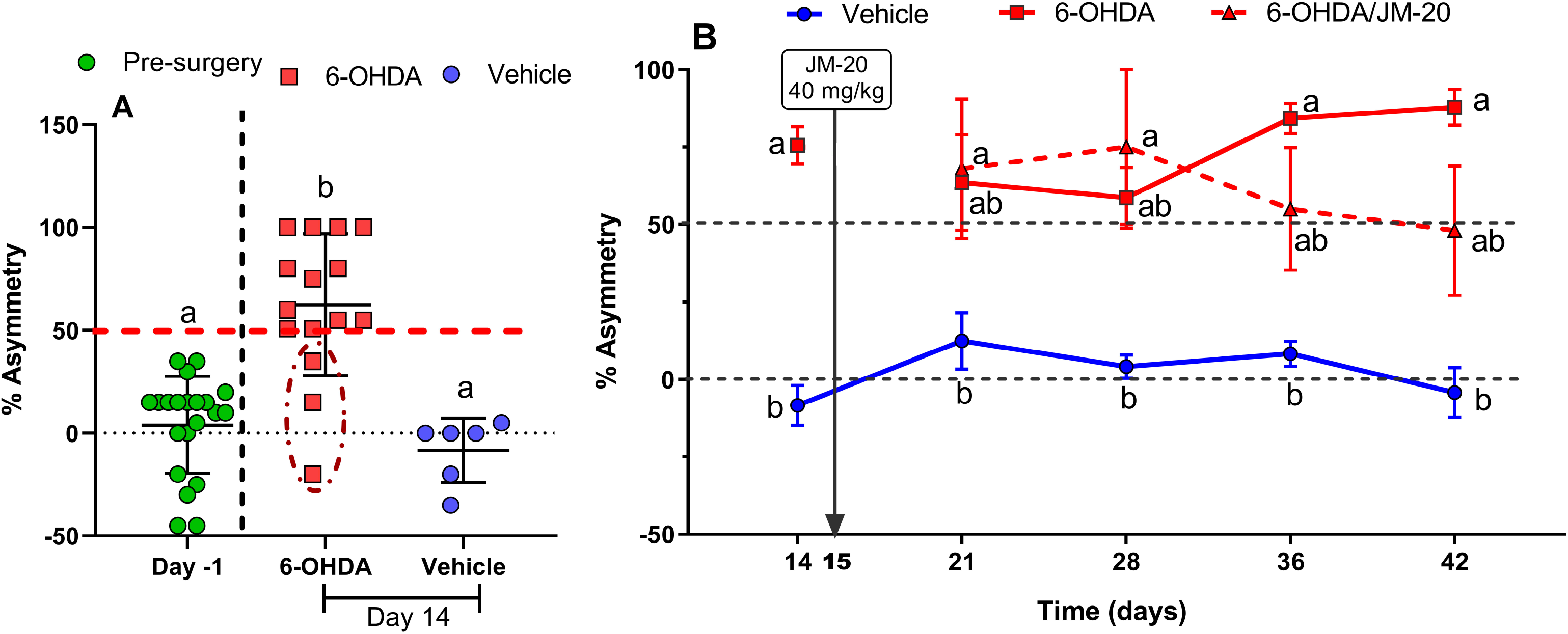
Protective properties of JM-20 against 6-OHDA-induced motor impairment evaluated in the cylinder test. (**A**) Evaluation on day -1, pre-surgery, reveals that all animals presented values of asymmetry between limbs below 50%. Intranigral injection of 6-OHDA causes a significant increase in percentage asymmetry (asymmetry≥50%) evaluated in day 14, in the majority of experimental subjects (12/15). The dashed line indicates subjects who did not show percent asymmetry above the cutoff value. Vehicle injection did not significantly modify the percentages of asymmetry with respect to untreated controls. The graph is presented as individual values ± SEM for each group. Different letters represent statistically significant differences, one-way ANOVA, followed by Tukey’s multiple comparison post-hoc test, (p < 0.0001). (**B**) The administration of JM-20 (40 mg/kg) 15 days post-surgery decreases the percentage of asymmetry in the use of the forelimbs of the treated animals. It is more evident from day 36 post-surgery. The graph is presented as mean values ± SEM of each group. Different letters represent statistically significant differences, two-way ANOVA, followed by Tukey’s multiple comparison post-hoc test, (p= 0.0006). Experimental groups: vehicle (n = 6); 6-OHDA (n = 7) and 6-OHDA/JM-20 (40 mg/kg, n = 5).

On day 15, those animals that presented asymmetry values ≥ 50% were selected and randomly assigned to be included in the groups: 6-OHDA without treatment (6-OHDA + CMC) (n = 7) or 6-OHDA + JM-20 (40 mg/kg) (n = 5). Of the 15 animals lesioned with 6-OHDA, only 12 met the inclusion criteria (asymmetry values ≥ 50%, for 80% effectiveness of stereotaxic surgery). Therefore, the remaining three were excluded from the experiment (**Fig. 1A**).

Twenty-one days after surgery, two-way repeated ANOVA (treatments × time as repeated measures) showed a significant interaction between time (days) and treatment (F (6, 45) = 4.902, p= 0.0006) on asymmetry evaluation of three evaluated groups (vehicle and 6-OHDA) in the cylinder test. 6-OHDA lesioned rats, presenting a high percentage of asymmetry as compared to the control group (vehicle) (**Fig. 1B**).

The high levels of asymmetry percentage were maintained in 6-OHDA lesioned groups during the 42 days in which they were evaluated in the cylinder test. On day 21 and 28, the 6-OHDA group (6-OHDA + CMC) without treatment presented elevated asymmetry values, being statistically different with respect to the control vehicle (p < 0.05 and 0.01, respectively) (**Fig. 1B**). In the evaluation of days 36 and 42, a worsening in the motor performance in the cylinder test was observed in experimental subjects from the 6-OHDA group (6-OHDA + CMC) with respect to control vehicle group (p < 0.0001) (**Fig. 1B**). However, the 6-OHDA-damaged animals that received JM-20 treatment (40 mg/kg) showed no significant differences from day 21 to the end of the experiment (day 42) with respect to either of the other two groups: 6-OHDA+CMC or vehicle (**Fig. 1B**). In the evaluation of days 36 and 42, a behavioral improvement in motor performance (lower asymmetry values) was observed in those animals that received the treatment (6-OHDA + JM-20) (**Fig. 1B**).

In order to evaluate the 6-OHDA-induced alterations in brain redox statuses and the impact of JM-20 treatment, T-SH groups and MDA levels were assessed in the SNpc and ST. T-SH and MDA levels were determined 42 days after surgery. As shown in **Figure 2**, the 6-OHDA-injured group significantly reduced T-SH levels in the SNpc (p < 0.05) (**Fig. 2 C**), but not in the ST (**Fig. 2 A**), and increased MDA levels in both regions studied (p < 0.001) (**Figs. 2 B, D**), compared to the vehicle control group. Treatment with JM-20 (40 mg/kg), in the damaged animals, decreased the MDA concentration in both regions (p < 0.001). In the case of T-SH levels, there was an increase in the concentration in both regions, but it was not statistically different from the untreated damaged animals.

**Figure 2.**
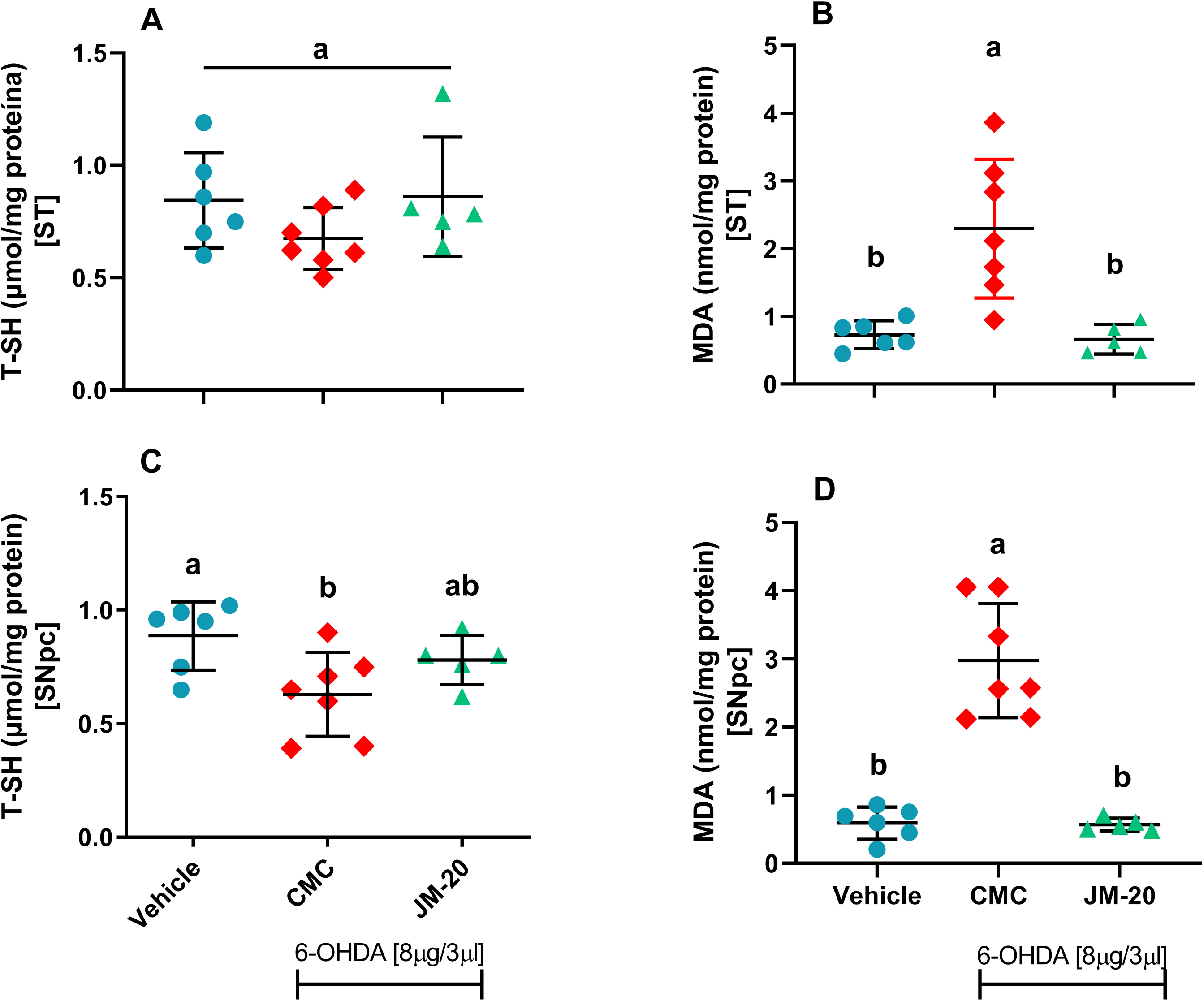
JM-20 improves the redox status of the SNpc and the ST in 6-OHDA lesioned rats. The figure shows the effect on the concentration of T-SH (**A and C**) and MDA (**B and D**) in the ST (**A-B**) and SNpc (**C-D**) of hemiparkinsonian (6-OHDA) animals and controls. Administration with JM-20 in hemiparkinsonian animals showed a decrease in MDA concentration in both the ST (**B**) and SNpc (**D**). No significant differences were found in the concentration of T-SH at the level of the ST (**A**) and SNpc (**C**) among the three experimental groups evaluated. In all cases, graphs are presented as individual values ± SEM for each group. Different letters represent statistically significant differences, one-way ANOVA, followed by Tukey’s multiple comparison *post-hoc* test, (p < 0.05). Experimental groups: vehicle (n = 6); 6-OHDA (n = 7) and 6-OHDA/JM-20 (40 mg/kg, n = 5).

## 4. Discussion

Several authors have reported that out of the motor tests, the cylinder test is one of the most sensitive and reliable for detecting the loss of dopaminergic neurons in the SNpc and one of the most consistent for detecting sensorimotor alterations in the 6-OHDA model in rodents ^21-23^. Likewise, Sauer and Oertel have reported that the injection of 6-OHDA into the SNpc caused progressive neurodegeneration, which started 1-2 weeks after the lesion and lasted 8-16 weeks after surgery ^24^. On the other hand, and aligned with our results, the group of Xiuping Sun et al. demonstrated that animals damaged with 6-OHDA and evaluated in the cylinder test exhibited high asymmetry values, which extended up to day 42 after surgery. They also reported a positive correlation between the degeneration of dopaminergic neurons (TH+) in the animals with the motor impairment observed in the cylinder test, as well as in other tests performed by the authors ^23^. Another group, in addition to finding the behavioral alterations typical of the model, found a significant redox imbalance, similar to our results. A decrease in glutathione levels was noted, as well as an increase in MDA and nitric oxide levels, and an increase in the production of reactive oxygen species 37 and 42 days after 6-OHDA injection, respectively ^25^.

There have been previous studies conducted by our group demonstrating different neuroprotective mechanisms of JM-20, which can prevent the loss of brain function in any neurodegenerative disease, particularly PD. Previous studies show that this molecule: I) presents strong antioxidant properties; II) modulates the glutamatergic system; III) preserves the mitochondrial functionality; IV) has anti-inflammatory and anti-apoptotic effects and V) prevents rotenone-induced motor damage, body weight loss and mortality, improving the redox state and inhibiting spontaneous mitochondrial swelling and membrane potential dissipation in the brain mitochondria of rats isolated after rotenone intoxication ^2-6,9,13,26^. In the authorś opinion, all these molecular mechanisms of the JM-20 that have been previously reported could explain the *in vivo* neurorestorative properties observed in the 6-OHDA preclinical models of PD, described in this work.

## Funding statement

This work was partially supported by SCIENCE AND INNOVATION FINANCIAL FUND (FONCI-Cuba) project 460020.

## Conflict of Interests statement

The authors declare that they have no conflict of interest.

## Availability of data and material

All data generated or analyzed during this study are included in this published article.

## Author contributions

(1) Research project: A. Conception, B. Organization, C. Execution; (2) Statistical Analysis: A. Design, B. Execution, C. Review and Critique; (3) Manuscript: A. Writing of the first draft, B. Review and Critique.

Luis Arturo Fonseca-Fonseca: 1A, 1B, 1C, 2A, 2B, 2C, 3A, 3B

Laura Reina Taño Portuondo: 1B, 1C, 2A, 2B, 2C, 3A, 3B

Jeney Ramírez-Sánchez: 1C, 2A, 2B, 3B

Abel Mondelo Rodríguez: 1B, 1C

Víctor Diogenes Amaral da Silva: 1A, 1B, 2A, 2B, 2C, 3A, 3B

Silvia Lima Costa: 1A, 1B, 2C, 3B

Yanier Núñez-Figueredo: 1A, 1B, 1C, 2A, 2B, 2C, 3A, 3B

## Notes

https://zenodo.org/records/13131465

### Competing Interest Statement

The authors have declared no competing interest.

## References

1 Nuñez-Figueredo, Y. et al. Multi-targeting effects of a new synthetic molecule (JM-20) in experimental models of cerebral ischemia. Pharmacological Reports 70, 699–704, doi:10.1016/j.pharep.2018.02.013 (2018).

2 Nunez-Figueredo, Y. et al. A novel multi-target ligand (JM-20) protects mitochondrial integrity, inhibits brain excitatory amino acid release and reduces cerebral ischemia injury in vitro and in vivo. Neuropharmacology 85, 517–527, doi:10.1016/j.neuropharm.2014.06.009 (2014).

3 Nunez-Figueredo, Y. et al. JM-20, a novel benzodiazepine-dihydropyridine hybrid molecule, protects mitochondria and prevents ischemic insult-mediated neural cell death in vitro. European journal of pharmacology 726, 57–65, doi:10.1016/j.ejphar.2014.01.021 (2014).

4 Ramirez-Sanchez, J. et al. Neuroprotection by JM-20 against oxygen-glucose deprivation in rat hippocampal slices: Involvement of the Akt/GSK-3beta pathway. Neurochemistry international 90, 215–223, doi:10.1016/j.neuint.2015.09.003 (2015).

5 Nunez-Figueredo, Y. et al. Antioxidant effects of JM-20 on rat brain mitochondria and synaptosomes: mitoprotection against Ca(2)(+)-induced mitochondrial impairment. Brain research bulletin 109, 68–76, doi:10.1016/j.brainresbull.2014.10.001 (2014).

6 Nunez-Figueredo, Y. et al. The effects of JM-20 on the glutamatergic system in synaptic vesicles, synaptosomes and neural cells cultured from rat brain. Neurochemistry international 81, 41–47, doi:10.1016/j.neuint.2015.01.006 (2015).

7 Nunez-Figueredo, Y. et al. Therapeutic potential of the novel hybrid molecule JM-20 against focal cortical ischemia in rats. J Pharm Pharmacogn Res 4, 153–158 (2016).

8 Nuñez-Figueredo, Y. et al. Multi-targeting effects of a new synthetic molecule (JM-20) in experimental models of cerebral ischemia. Pharmacological Reports 70, doi:10.1016/j.pharep.2018.02.013 (2018).

9 Ramírez-Sánchez, J. et al. JM-20 Treatment After MCAO Reduced Astrocyte Reactivity and Neuronal Death on Peri-infarct Regions of the Rat Brain. Molecular neurobiology 56, 502–512, doi:10.1007/s12035-018-1087-8 (2019).

10 Wong-Guerra, M. et al. JM-20 protects memory acquisition and consolidation on scopolamine model of cognitive impairment. Neurological research 41, 385–398, doi:10.1080/01616412.2019.1573285 (2019).

11 Wong-Guerra, M. et al. JM-20 treatment prevents neuronal damage and memory impairment induced by aluminum chloride in rats. Neurotoxicology 87, 70–85, doi:10.1016/j.neuro.2021.08.017 (2021).

12 Furtado, A. B. V. et al. JM-20 Treatment After Mild Traumatic Brain Injury Reduces Glial Cell Pro-inflammatory Signaling and Behavioral and Cognitive Deficits by Increasing Neurotrophin Expression. Molecular neurobiology 58, 4615–4627, doi:10.1007/s12035-021-02436-4 (2021).

13 Fonseca-Fonseca, L. A. et al. JM-20, a novel hybrid molecule, protects against rotenone-induced neurotoxicity in experimental model of Parkinson’s disease. Neuroscience letters 690, 29–35, doi:10.1016/j.neulet.2018.10.008 (2019).

14 Fonseca-Fonseca, L. A. et al. JM-20 protects against 6-hydroxydopamine-induced neurotoxicity in models of Parkinson’s disease: Mitochondrial protection and antioxidant properties. Neurotoxicology 82, 89–98, doi:10.1016/j.neuro.2020.11.005 (2021).

15 Santos, C. C. et al. JM-20, a Benzodiazepine-Dihydropyridine Hybrid Molecule, Inhibits the Formation of Alpha-Synuclein-Aggregated Species. Neurotoxicity research 40, 2135–2147, doi:10.1007/s12640-022-00559-7 (2022).

16 Woodlee, M. T., Kane, J. R., Chang, J., Cormack, L. K. & Schallert, T. Enhanced function in the good forelimb of hemi-parkinson rats: compensatory adaptation for contralateral postural instability? Experimental neurology 211, 511–517, doi:10.1016/j.expneurol.2008.02.024 (2008).

17 Boix, J., Padel, T. & Paul, G. A partial lesion model of Parkinson’s disease in mice--characterization of a 6-OHDA-induced medial forebrain bundle lesion. Behavioural brain research 284, 196–206, doi:10.1016/j.bbr.2015.01.053 (2015).

18 Su, R. J. et al. Time-course behavioral features are correlated with Parkinson’s diseaselZIassociated pathology in a 6-hydroxydopamine hemiparkinsonian rat model. Molecular medicine reports 17, 3356–3363, doi:10.3892/mmr.2017.8277 (2018).

19 O’Brien, J. A. & Austin, P. J. Effect of Photobiomodulation in Rescuing Lipopolysaccharide-Induced Dopaminergic Cell Loss in the Male Sprague-Dawley Rat. Biomolecules 9, doi:10.3390/biom9080381 (2019).

20 Wong-Guerra, M., Jimenez-Martin, J. & Pardo-Andreu, G. L. Mitochondrial involvement in memory impairment induced by scopolamine in rats. 39, 649–659, doi:10.1080/01616412.2017.1312775 (2017).

21 Iancu, R., Mohapel, P., Brundin, P. & Paul, G. Behavioral characterization of a unilateral 6-OHDA-lesion model of Parkinson’s disease in mice. Behavioural brain research 162, 1–10, doi:10.1016/j.bbr.2005.02.023 (2005).

22 Glajch, K. E., Fleming, S. M., Surmeier, D. J. & Osten, P. Sensorimotor assessment of the unilateral 6-hydroxydopamine mouse model of Parkinson’s disease. Behavioural brain research 230, 309–316, doi:10.1016/j.bbr.2011.12.007 (2012).

23 Sun, X. et al. Longitudinal assessment of motor function following the unilateral intrastriatal 6-hydroxydopamine lesion model in mice. Frontiers in Behavioral Neuroscience 16, 982218, doi:10.3389/fnbeh.2022.982218 (2022).

24 Sauer, H. & Oertel, W. H. Progressive degeneration of nigrostriatal dopamine neurons following intrastriatal terminal lesions with 6-hydroxydopamine: a combined retrograde tracing and immunocytochemical study in the rat. Neuroscience 59, 401–415, doi:10.1016/0306-4522(94)90605-x (1994).

25 Tiwari, P. C., Chaudhary, M. J., Pal, R. & Nath, R. Effects of mangiferin and its combination with nNOS inhibitor 7-nitro-indazole in 6-OHDA lesioned Parkinson’s disease rats. Fundamental & Clinical Pharmacology 36, 944–955, doi:10.1111/fcp.12817 (2022).

26 Figueredo, Y. N. et al. Characterization of the anxiolytic and sedative profile of JM-20: a novel benzodiazepine-dihydropyridine hybrid molecule. Neurological research 35, 804–812, doi:10.1179/1743132813Y.0000000216 (2013).

